# Replicating dynamic immune responses at single-cell resolution within a microfluidic human skin equivalent

**DOI:** 10.1101/2024.11.15.623786

**Authors:** Sarah A. Hindle, Holly Bachas Brook, Alexandra Chrysanthou, Emma S. Chambers, Matthew P. Caley, John T. Connelly

## Abstract

The intricate immunological functions of human skin involve the interplay between multiple different cell types as well as dynamic trafficking of leukocytes in and out the tissue, both of which are extremely challenging to replicate in vitro. To enable in vitro investigation of human skin immunology, we developed a microfluidic human skin equivalent (HSE) that supports the delivery of circulating immune cells via a vascular microchannel embedded within the dermis of a full-thickness construct. We demonstrated that stimulation of keratinocyte inflammation with lipopolysaccharide and nigericin promoted rapid monocyte recruitment out of the vascular channel and into the epidermal layer within 24 hours, followed by a second wave of monocyte migration into the dermis over a period of six days. Single-cell transcriptomic analysis of the tissue-resident and recruited cell populations revealed dynamic and cell-specific patterns of gene expression that were characteristic of acute activation and resolution of an inflammatory immune response.

Moreover, comparison of the gene signatures of the monocyte-derived cells to in vivo populations provided molecular level validation of the model and indicated a differentiation trajectory of the monocytes through to mature dermal macrophages. To extend the microfluidic platform to additional applications, we also modelled age-associated immune dysfunction by the inclusion of senescent fibroblasts, which promoted increased monocyte recruitment into the HSE, replicating previous in vivo human studies. Thus, the microfluidic HSE presented here replicates key aspects of dynamic inflammatory immune responses within the skin and represents a tractable experimental tool for interrogating mechanisms of human skin immunology.

## Introduction

Human skin is a major physical and immunological barrier between the body and the external environment, and it carries out its complex protective functions through crosstalk between tissue resident cells, circulating immune cells, and secondary lymphoid tissues. Skin resident immune cells include a diverse repertoire of macrophages, mast cells, dendritic cells, and lymphocytes that respond directly to foreign pathogens but also modulate the local tissue microenvironment^1^. In addition, immune cells continuously migrate in and out of the skin via the vasculature and lymphatics to control infections, maintain tissue homeostasis, and communicate with lymphoid tissues^1^. Regulation of immune cell activity within the skin also depends on communication with the resident keratinocytes, dermal fibroblasts, and blood vessels, which are all key players in the tissue’s overall immune function^1^.

Given the multicellular, multiorgan, and dynamic nature of skin immunology, research to date has relied heavily on mouse models to study these complex processes. However, there is growing appreciation of the physiologic differences between human and mouse skin^2,3^ and the ethical concerns regarding animal experimentation^4^. Human skin equivalents (HSE) and organotypic models have become valuable tools for replicating the three-dimensional (3D) structure of human skin and studying mechanisms of tissue development and homeostasis^5–7^. These models often consist of bi-layered dermal and epidermal compartments with fibroblasts and keratinocytes, but they typically lack more complex features, including vasculature, immune cells, nerves, and appendages.

Recent work has begun to build vascular structures into 3D skin models through the use of microfluidic channels^8–10^ or self-organising microvascular networks^11^, and some have likewise, incorporated tissue-resident immune cells, such as macrophages, T cells, and Langerhans cells^12–15^. However, fully vascularised and immuno-competent skin models that support the dynamic trafficking of immune cells still remain a significant challenge. In a couple recent studies, monocytes and neutrophils were successfully circulated through vascular compartments in microfluidic skin models^12,16,17^, suggesting that these types of cells can be incorporated into in vitro skin models, but it has yet to be determined whether these types of approaches can replicate all the essential processes involved in the immune response to infection or skin inflammation.

Monocytes in particular play key roles in the innate response to infection and have the potential to differentiate into tissue resident myeloid cells, to replenish the pool of macrophages, dermal dendritic cells, and Langerhans cells and maintain skin homeostasis^18,19^. In addition, dysfunctional monocyte responses have been implicated in a range of inflammatory skin diseases and in the age-associated decline in immunity^20,21^. However, the underlying molecular mechanisms and intercellular signalling networks that direct monocyte fate and function are not fully understood, and advanced human skin models that support dynamic trafficking of immune cells could open up new opportunities to dissect the roles of monocytes in human skin health and disease.

In this study, we constructed microfluidic HSEs that consisted of an endothelial-lined microchannel, running through a 3D dermal matrix embedded with fibroblasts, and a layer of epidermal keratinocytes on the surface. Human monocytes were delivered to the tissue via the vascular microchannel and specifically recruited into the HSE upon inflammatory activation of the keratinocytes. We further leveraged live imaging and single-cell transcriptomics to characterise the cellular and molecular dynamics of this response. Normal monocyte responses could also be perturbed through the addition senescent fibroblasts to the HSE, thereby mimicking age-related immune dysfunction. Together, these studies establish a novel model and methodology for emulating acute and chronic immune responses in the skin and provide molecular-level benchmarking and insights into the underlying regulatory mechanisms.

## Results

### Fabrication of 3D skin equivalents with microfluidic vascular channels

To construct vascularised HSEs that support fluidic delivery of immune cells, we engineered dermal matrices with microfluidic channels using 3D printing with a sacrificial material. The overall fabrication process (Figure 1a,b) involved first 3D printing a silicone frame with inlet and outlet ports to house the microfluidic HSE. A fibrin gel embedded with primary human dermal fibroblasts was then cast in the bottom of the frame, and the sacrificial template for the microchannel was created by 3D printing a line of gelatin on the fibrin gel between the two ports. The gelatin template remained solid at room temperature, and a second fibroblast-embedded fibrin gel was cast over the top. The construct was then incubated at 37°C to melt the gelatin, and the liquified gelatin was flushed out of the channel with warm medium, leaving behind a hollow microchannel running through the 3D dermal matrix.

**Figure 1:**
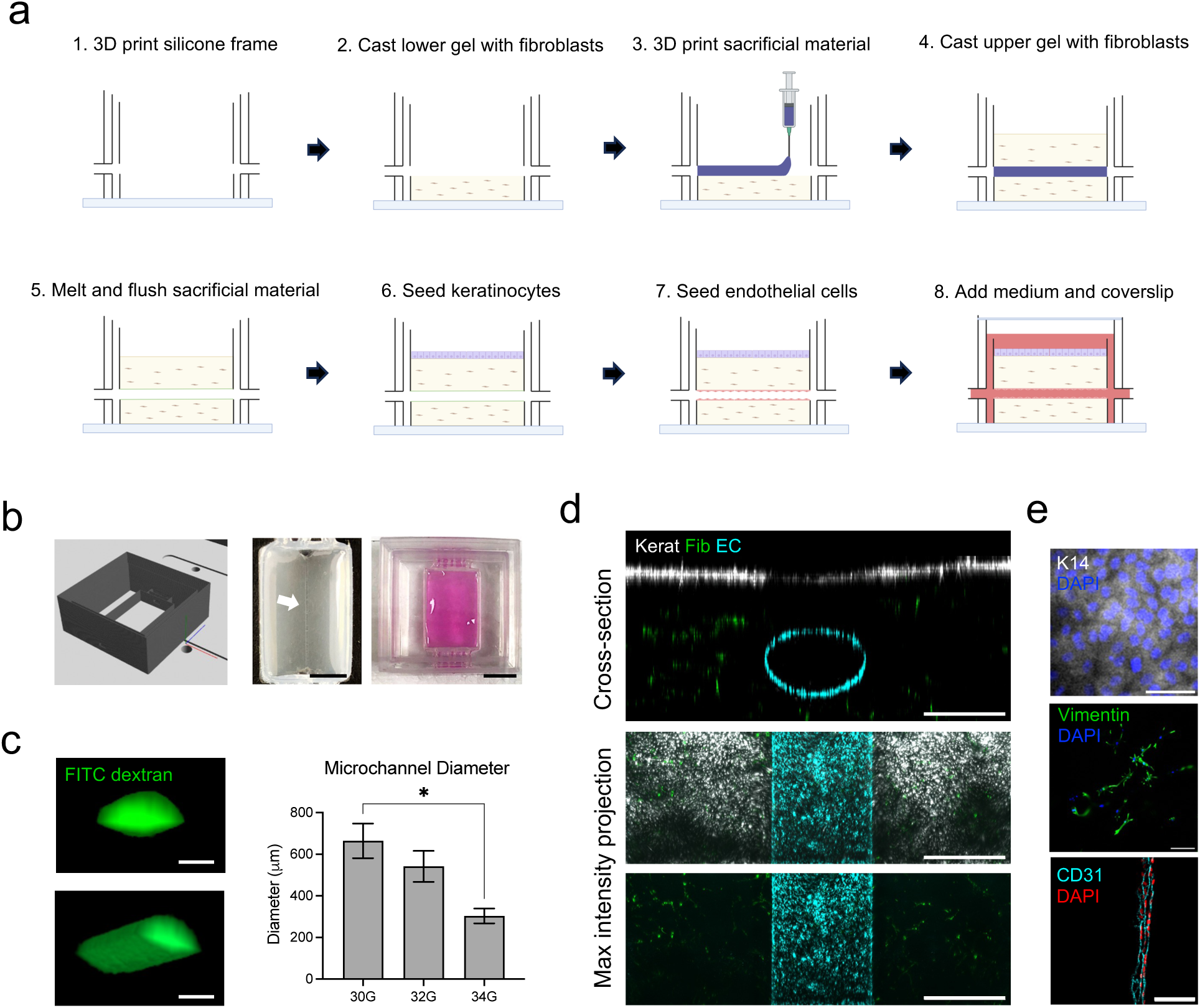
Fabrication of a microfluidic human skin equivalent. (a) Schematic of the fabrication process for constructing the microfluidic human skin equivalent (HSE) using 3D printing with sacrificial materials. (b) Computer-aided design image of the silicone frame (left), photo of the HSE with a gelatin template for the microchannel (middle), and photo of the fully formed HSE with culture medium in the inner chamber (right). Scale bar = 5 mm. (c) Cross-section and 3D rendering of microchannels filled with 40 kDa FITC-dextran imaged via confocal microscopy, and quantification of average channel diameter in HSEs fabricated with varying printhead needle sizes. Data represent the mean of N=3 experiments, *p<0.05, ANOVA. Scale bar = 200 μm. (d) Cross-section and maximum intensity images of HSEs containing N/TERT keratinocytes (Kerat), primary human dermal fibroblasts (Fib), and human umbilical cord vascular endothelial cells (EC) each labelled with different fluorescently tagged membrane dyes and imaged via confocal microscopy. Scale bar = 500 μm. (e) Representative immunofluorescence images of cell-specific markers, including keratin 14 for keratinocytes on the surface, vimentin for fibroblasts within the dermal matrix, and CD31 for ECs along the wall of the microchannel. Scale bar = 100 μm.

We confirmed the formation and geometry of the microchannels by injecting FITC-dextran (40 kDa) into the channel and imaging by confocal microscopy. The microchannels had an oval cross-section due to slight spreading of the gelatin template, and the average width of the channel could be adjusted between approximately 300 μm and 700 μm, depending on the size of the print head needle (Figure 1c). For all subsequent studies, a 32G needle was used to produce 500 μm channels.

N/TERT keratinocytes were then seeded on the surface of the construct to form an epidermal layer, and human umbilical vein endothelial cells (ECs) were injected into the microchannel to form an endothelial lining and mimic the vasculature. To reduce the fabrication time and simplify the model, the keratinocytes were cultured as an epithelial monolayer submerged in medium, rather than a fully stratified epidermal layer, which is feasible within this platform (Figure S1a). To visualise the three different cell types within the model, fibroblasts, keratinocytes, and ECs were each labelled with fluorescent membrane dyes prior to seeding into the HSE. Confocal imaging of the microfluidic HSE confirmed that the ECs formed a confluent endothelial lining along the length of the microchannel with expression of CD31 at the cell-cell junctions (Figure 1d-e). Fibroblasts displayed an elongated morphology throughout the 3D dermal matrix and stained positively for vimentin (Figure 1d-e). Keratinocytes formed a confluent epithelial layer on the surface and stained positively for keratin 14 (Figure 1d-e), and these co-cultures could be maintained for at least one week in the EC medium, EGM2. This fabrication strategy therefore established a robust method for building 3D HSEs with microfluidic vascular channels within the dermis.

### Activation of inflammation and monocyte recruitment

To investigate monocyte trafficking within the microfluidic HSE, inflammatory signalling was first activated in the keratinocytes to promote monocyte recruitment from the vascular channel and into the HSE (Figure 2a). The epidermal surface was stimulated by a two-step treatment to prime toll-like receptor signalling with 500 ng/ml lipopolysaccharide and then activate the NLRP3 (NOD- LRR- and pyrin domain-containing protein 3) inflammasome with 10 μM nigericin (LPS/nig)^22,23^. This protocol promoted secretion of multiple chemokines and inflammatory cytokines, including CXCL12, IL1-α, IL-18, and IL-8 (Figure 2b). Monocytes labelled with fluorescent membrane dyes were then injected into the microchannel and their migration was tracked through the HSE by confocal microscopy.

**Figure 2:**
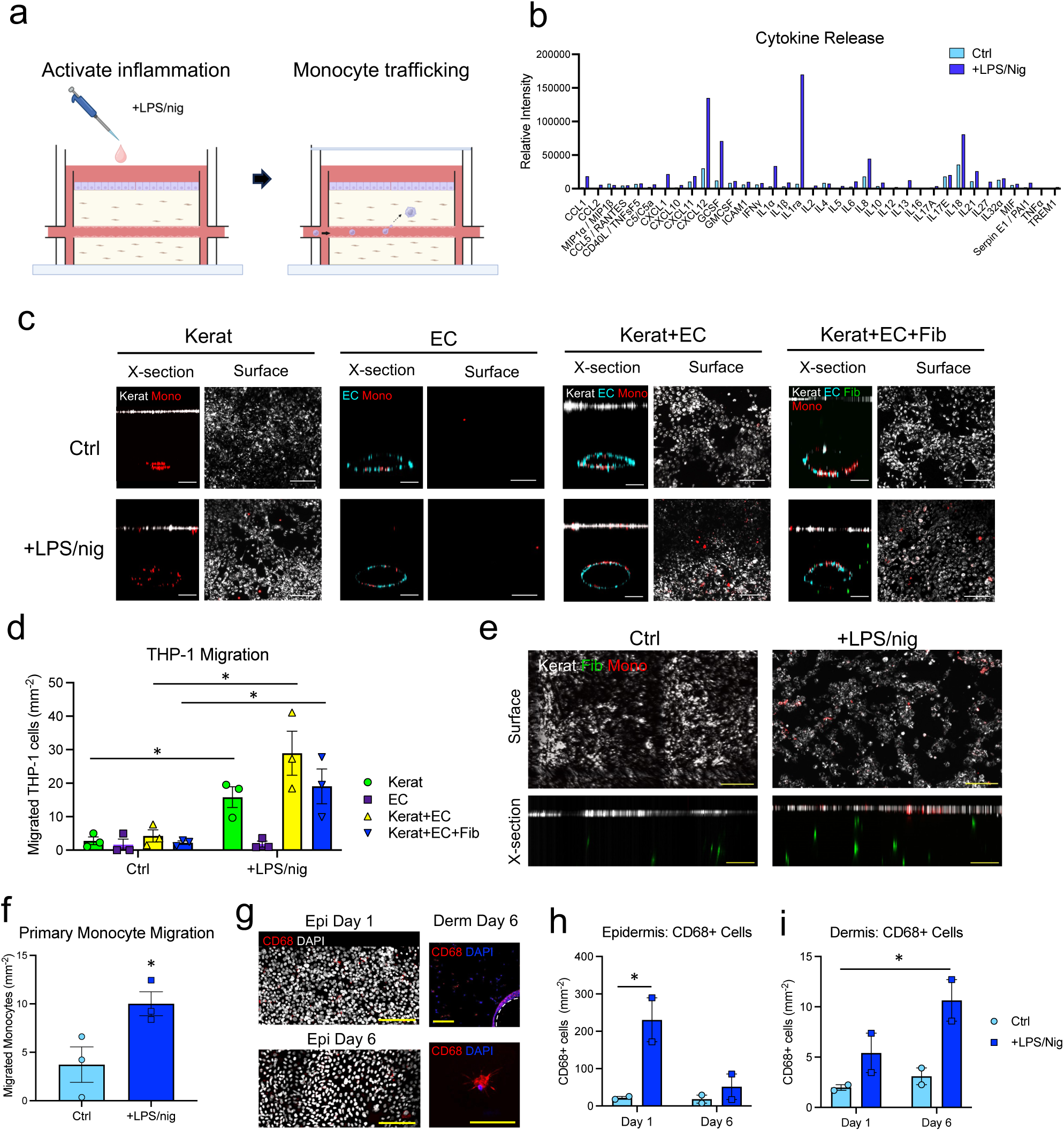
Activation and analysis of monocyte recruitment. (a) Schematic of activation of inflammation by treatment of the HSEs with 500 ng/ml LPS for 4 h, followed by 10 μM nigericin (LPS/nig) for 1 h, and then injecting monocytes into the vascular microchannel. (b) Quantification of the cytokine profile in the medium of N/TERT keratinocytes 24 h after treatment with control (Ctrl; untreated) medium or two-step stimulation with LPS/nig. Data represent the average intensity (arbitrary units) for duplicate spots from the cytokine array. (c) Representative confocal microscopy images of cross-section and surface views of HSEs constructed with different combinations of fluorescently tagged Kerats, Fibs, and ECs, treated with LPS/nig, and injected with fluorescently tagged THP-1 monocytes. HSEs were imaged 24 h after monocyte injection. Scale bart = 200 μm. (d) Quantification of THP-1 monocyte density in the epidermal layer at 24 h. Data represent the mean ± SEM of N=3 experiments, *p<0.05 compared to Ctrl, ANOVA. (e) Representative confocal microscopy images of cross-section and surface views of HSEs treated with LPS/nig or Ctrl medium and injected with fluorescent primary human monocytes (CD14+). HSEs were imaged 24 h after monocyte injection. Scale bar = 200 μm. (f) Quantification of primary monocyte density in the epidermal layer at 24 h. Data represent the mean ± SEM of N=3 experiments, *p<0.05 compared to Ctrl, ANOVA. (g) Immunofluorescence images of CD68 expression in cells in the epidermal (surface view) and dermal (cross-section view) compartments of the HSEs at days 1 and 6 after LPS/nig treatment. Scale bar = 100 μm. (h) Quantification of CD68+ cell density in the epidermal layer and (i) dermis for Ctrl and LPS/nig treated HSEs at days 1 and 6. Data represent the mean ± range of N=2 experiments, *p<0.05 compared to Ctrl day 1, ANOVA.

In initial studies, simplified models containing only keratinocytes and the THP-1 monocytic cell line demonstrated that treatment with LPS/nig stimulated monocyte migration out of the microchannel within 3 h, and by 24 h there was significant migration of monocytes up to the epidermal layer (Figure S1b,c). Nearly all migrating monocytes were found in the epidermal layer with only an occasional cell observed in the dermis. Migration depended on signals from keratinocytes as minimal migration was observed in the monocyte only cultures (Figure S1b,c).

Activation of THP-1 monocyte migration was also observed in cultures containing keratinocytes, ECs, and fibroblasts, where stimulation with LPS/nig significantly increased monocyte migration up to the epidermal layer of the HSE after 24 h. Similar levels of migration were observed in all the models with keratinocytes, but there was minimal migration in the EC only cultures (Figure 2c-d). These results indicated that inflammatory signals produced by the keratinocytes were the main drivers of monocyte recruitment into the microfluidic HSE.

We next confirmed that activation of inflammation in the microfluidic HSE also stimulated migration of primary human monocytes. CD14+ human monocytes were isolated from the blood of healthy volunteers and injected into the microchannels, and a significant increase in monocyte migration up to the epidermal layer was observed in LPS/nig treated models at 24h (Figure 2e,f). Like the THP-1 cells, only a few monocytes were found within the dermis at this time point. Monocyte fate within these models was further explored by immunofluorescence staining at early (24 h) and later (day 6) time points. Cells expressing the pan-macrophage marker CD68 were observed in the microchannel, dermis, and epidermis at both days 1 and 6 with overall higher numbers in the epidermis and in the LPS/nig treated cultures (Figure 2g-i). The density of CD68+ cells in the epidermis decreased from day 1 to day 6, while the density in the dermis continued to increase (Figure 2h-i), suggesting differing kinetics of migration into these two compartments. Given the presence of monocytes in the epidermal layer, we also examined the Langerhans cell marker, CD207, but observed no positive cells (data not shown).

Together, these experiments demonstrated that stimulation of the microfluidic HSE with LPS/nig activated a potent inflammatory response that promoted monocyte migration into the dermal and epidermal compartments. Expression of CD68 at days 1 and 6 further suggested that the monocytes rapidly began differentiating along a macrophage-like lineage within the HSE model.

### Transcriptomic analysis of dynamic inflammatory responses in tissue-resident cells

To gain a deeper molecular understanding into monocyte fate and the intercellular regulatory networks within the system, we performed single-cell transcriptomic analysis of cells within the microfluidic HSEs following inflammatory activation with LPS/nig. Constructs containing keratinocytes, fibroblasts, ECs and primary monocytes were analysed by single-cell RNA-sequencing (10X Genomics), and approximately 6,000-8,000 cells per condition were successfully sequenced at days 1 and 6 following LPS/nig stimulation alongside untreated controls. Initial UMAP visualisation of all cells identified 20 distinct clusters, which were classified based on cell type-specific markers for each major cell type in the model (Figure 3a-b; Supplementary Data S1). This analysis identified 9 clusters of keratinocytes, 7 clusters of fibroblasts, 2 clusters of ECs, and 2 clusters of monocytes, confirming that all four cell types were present within the data set.

**Figure 3:**
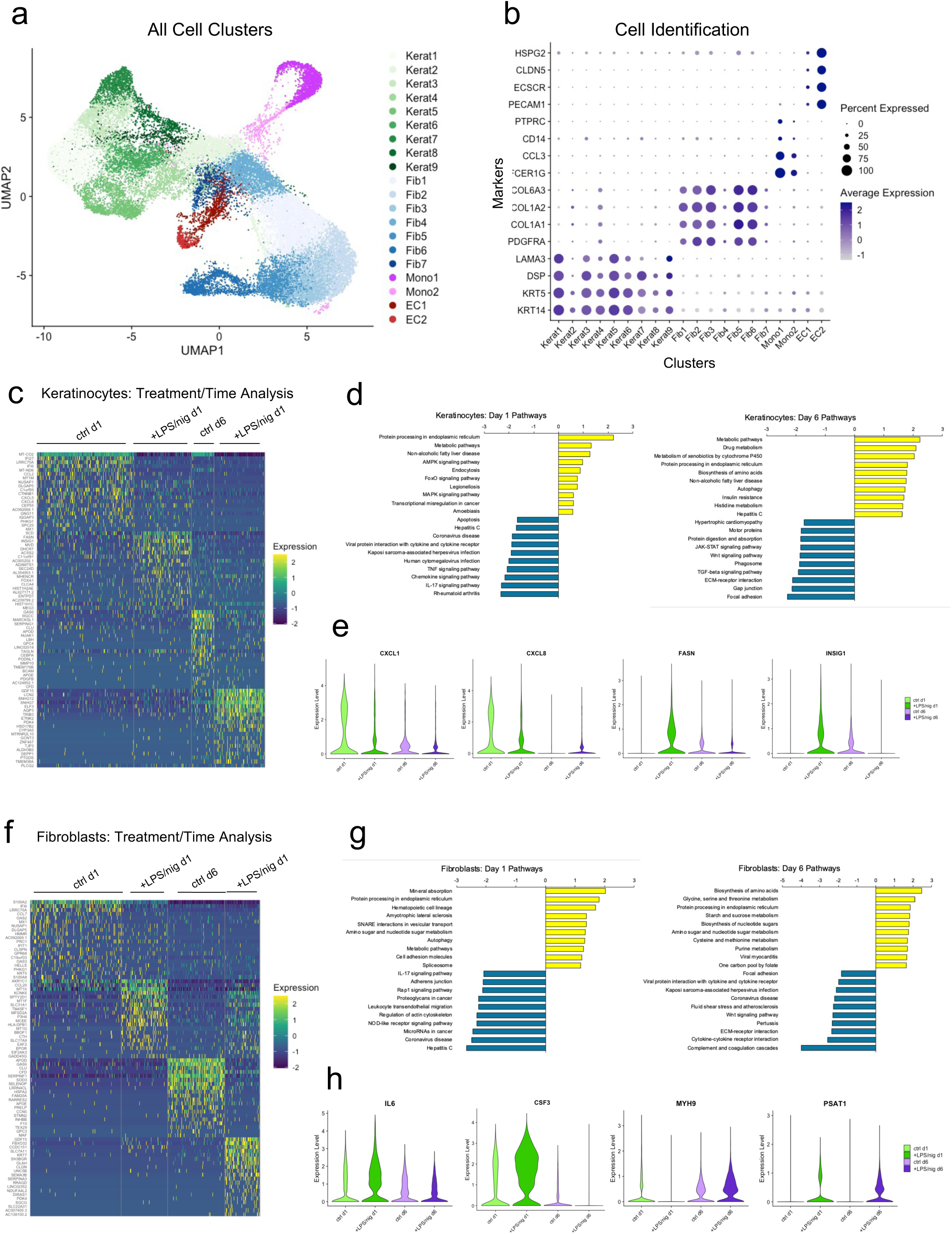
Single-cell transcriptomic analysis of inflammation dynamics within the microfluidic HSE. (a) Integrated uniform manifold approximation and projection (UMAP) visualisation of single-cell RNA-seq analysis within the microfluidic HSE model. Data represent all cells that passed quality control criteria from ctrl (untreated) and LPS/nig treated HSEs at days 1 and 6 (N=3 HSEs per condition). The cell type for each cluster was identified using conserved markers for keratinocytes, fibroblasts, ECs, and monocytes (b) Dot plot showing the percentage of cells and average expression level for the cell type specific markers within each cluster. (c) Heatmap of the top 20 differentially expressed genes (DEGs; Log2FC > 1; Padj < 0.05) between experimental conditions for all keratinocytes. (d) Gene set enrichment analysis (GSEA; Webgestalt) of keratinocyte pathways up and down regulated by LPS/nig at days 1 and 6. Data represent pathways with the top 10 highest and lowest relative enrichment scores. (e) Violin plots showing the relative expression and distribution of selected genes (*CXCL1, CXCL8 FASN, INSIG1*) in all keratinocytes grouped by experimental conditions. (f) Heatmap of top 20 differentially expressed genes (DEGs; Log2FC > 1; Padj < 0.05) between experimental conditions for all fibroblasts. (g) Gene set enrichment analysis (GSEA; Webgestalt) of fibroblast pathways up and down regulated by LPS/nig at days 1 and 6. Data represent pathways with the top 10 highest and lowest relative enrichment scores. (h) Violin plots showing the relative expression and distribution of selected genes (*IL6, CSF3, MYH9, PSAT1*) in all fibroblasts grouped by experimental conditions.

We then analysed the transcriptomic signatures of the tissue-resident cell types to first understand how the local microenvironment of the HSE changed in response to inflammatory activation with LPS/nig. Within the keratinocytes, cell clusters segregated primarily based on their differentiation state and expression of inflammatory markers (Figure S2a,c). The keratinocyte clusters also segregated according to treatment and time, with the LPS/nig treated cells aligning with more differentiated clusters expressing *KRT10* and clusters with high expression *IL18* (Figure S2b-d). By contrast, the day 1 control cells were found mostly in clusters with low levels of *KRT10* and *IL18* and high levels of the chemokine *CXCL1*, but by day 6, control cells shifted to *IL18* high / *CXCL1* low clusters (Figure S2b-d). We performed differential gene expression (DEG) and pathway analysis across all keratinocyte populations and identified distinct patterns of expression and enriched pathways that varied based on both treatment and time (Figure 3c; Supplementary Data S2). At day 1, we observed an upregulation of genes related to protein and lipid metabolism pathways, such as *FASN* and *INSIG1*, in the LPS/nig treated cells compared to untreated controls (Figure 3d,e). There was also an unexpected downregulation of genes associated with viral infection and several inflammatory pathways, including TNF, chemokine, and IL-17 signalling, (e.g. *CXCL1* and *CXCL8*). At day 6, additional metabolic pathways were upregulated in the LPS/nig treated keratinocytes, while pathways related to Wnt, TGF-β, and ECM adhesion were downregulated (Figure 3d,e).

Together, these data demonstrated a dynamic transcriptional response within the keratinocytes following stimulation with LPS/nig, and this response was characterised by the upregulation of *IL18* alongside several metabolic pathways. Interestingly, the early downregulation of inflammatory pathways may reflect initial feedback mechanisms within the keratinocytes to resolve the inflammatory response initiated by LPS/nig treatment.

Similar to the keratinocytes, we also analysed the response of the fibroblasts to LPS/nig treatment. Here, the fibroblast clusters segregated by expression of cytokine and chemokine genes, such as *CCL2* and *CSF3*, and by genes associated with the ECM and contractility, such as *COL6A1* and *MYL9* (Figure S3a-d). The fibroblast clusters also segregated by treatment and time. Expression of *CCL2* and *CSF3* differentiated between control and LPS/nig treated fibroblasts at day 1, while most day 6 fibroblasts aligned with the clusters with high expression of ECM and contractility genes (Figure S3b-d). Further analysis of the DEGs and enriched pathways across all the fibroblasts revealed a transient upregulation of several key inflammatory cytokines, such as *IL6*, *CSF3*, and *IL1B*, at day 1 in the LPS/nig treated fibroblasts (Figure 3g,h; Figure S3d; Supplementary Data S2). Conversely, there was a transient downregulation of cytoskeletal genes, such as *MYH9* and *ACTB*, (Figure 3h; Figure S3d) in response to LPS/nig treatment. We also observed an upregulation of metabolic pathways and genes such as *PSAT1*, in the LPS/nig groups at both time points, and there was a downregulation of pathways associated with viral infections at both time points and a downregulation of the complement pathway, specifically at day 6 (Figure 3g,h). These data therefore demonstrated a dynamic transcriptional response within the fibroblast population upon stimulation with LPS/nig and provided evidence for both activation and resolution of inflammatory gene expression within this system.

A similar analysis of transcriptional responses was also performed for the ECs. Overall, there were fewer differentially expressed genes in the ECs, and the two cell clusters segregated mainly by time point (Figure S4a). However, notable pathways upregulated by LPS/nig included genes associated with viral infections, NF-kappa B signalling, and cell adhesion, and the temporal changes in *ICAM1*, *NFKB1*, and *VWF* confirm that ECs also display a dynamic inflammatory response to LPS/nig treatment (Figure S4b-f; Supplementary Data S2). Taken together, these findings indicated that each of the resident cell populations within the microfluidic HSE displays a heterogenous and dynamic transcriptional response to inflammatory activation of the system. Moreover, downregulation of inflammatory genes and pathways over the 6 day time course suggested that the inflammatory response of the tissue resident cells of the HSE began to resolve during this time period.

### Transcriptomic analysis of monocyte fate and regulatory pathways

We next performed a more detailed investigation of monocyte fate within the microfluidic HSE models and re-clustered the monocyte-derived populations to gain a more refined understanding of the different cell states. With this analysis, the monocytes separated into eight clusters, similar to previously reported levels of population heterogeneity in acute wound healing studies^24^, and the clusters aligned with different time points and treatment conditions (Figure 4a-c). Monocyte clusters 1 and 2 (Mo1 and Mo2) comprised mostly day 1 control cells, while day 1 LPS/nig treated cells were found mostly in the Mo3-6 clusters. The day 6 cells for both control and LPS/nig treated conditions aligned most closely with Mo7 and Mo8 (Figure 4c). To further assess the phenotype of the monocyte clusters we compared the top 100 DEGs for each cluster to published gene signatures for all myeloid cells found in human skin^25^. While Mo1-3 displayed little similarity to any of the myeloid populations, Mo4-6, which consisted mostly of day 1 LPS/nig treated cells, were most similar to monocyte-derived macrophages and dendritic cells (DCs) (Figure 4d; Supplementary Data S3). By contrast, the day 6 clusters were most similar to mature tissue resident cells. Mo7 was most similar to DC 2 and Langerhans cell (LC) 1 populations, and Mo8 displayed high similarity to macrophage 1 cells with expression of skin resident macrophage markers, such as *LYZ* and *CD163* (Figure 4d,e). Moreover, trajectory analysis using Monocle3 revealed that progression along pseudotime aligned with the differentiation state of the monocyte clusters (Figure 4f). Together, these results indicated that monocytes within the microfluidic HSE differentiated towards mature skin-resident myeloid cells. Stimulation of inflammation with LPS/nig promoted early differentiation into monocyte-derived DCs and macrophages, but by day 6, cells in both conditions transcriptionally resembled tissue-resident DCs, LCs, and macrophages.

**Figure 4:**
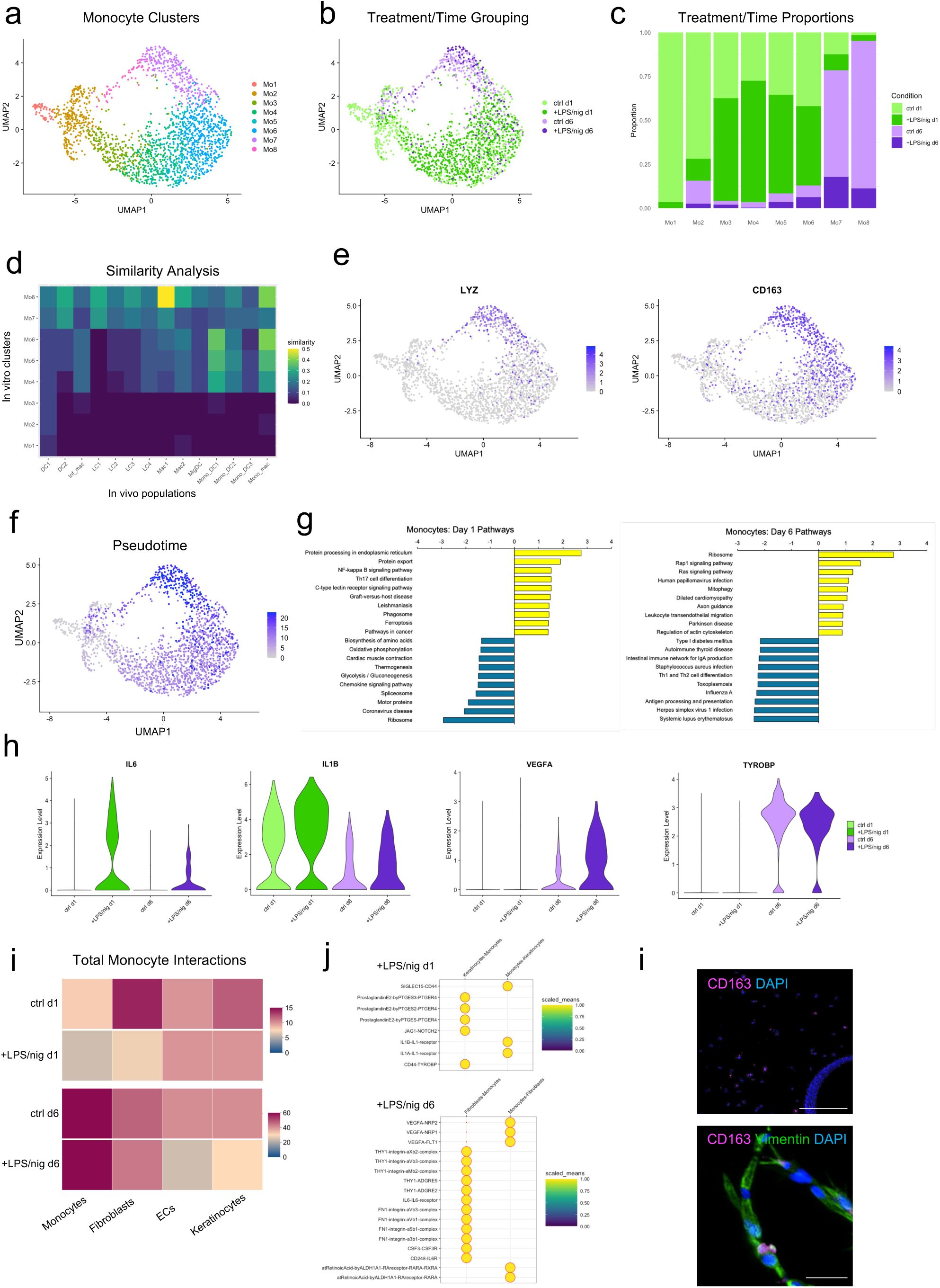
Single-cell analysis of monocyte fate and function within the microfluidic HSE. (a) UMAP visualisation of eight monocyte-derived populations after re-clustering all monocytes identified in the complete RNA-seq data set. (b) UMAP visualisation of monocyte clusters grouped by experimental condition. (c) Quantification of the proportion of each cluster comprised of monocytes from different experimental treatments. (d) Heatmap showing the relative level of similarity between the HSE monocyte clusters and myeloid populations present in human skin^25^. Similarity was calculated as the fraction of the signature genes for each human skin population present in the top 100 DEGs for each monocyte cluster from the HSEs. (e) Feature plot for relative expression of macrophage markers *LYZ* and *CD163* within the monocyte-derived populations. (f) Feature plot of relative changes in pseudotime across the monocyte populations calculated using Monocle3. (g) Gene set enrichment analysis (GSEA; Webgestalt) of monocyte pathways up and down regulated by LPS/nig at days 1 and 6. Data represent pathways with the top 10 highest and lowest relative enrichment scores. (h) Violin plots showing the relative expression and distribution of selected genes (*IL6, IL1B, VEGFA, TYROBP*) in all monocytes grouped by experimental conditions. (i) Heatmap of all predicted molecular interactions (CellPhoneDB) between monocytes and tissue resident cells across the different experimental conditions. (j) Dot plot of selected significant interactions between the monocytes and keratinocytes in HSEs treated with LPS/nig at day 1 and monocytes and fibroblasts in HSEs treated with LPS/nig at day 6. Intensity of the scaled means corresponds to normalised interaction scores. (k) Representative immunofluorescence images of CD163 and vimentin in the dermal compartment of day 6 HSEs treated with LPS/nig. Scale bars correspond to 200 μm (upper) and 50 μm (lower).

To understand the underlying signalling pathways involved in the monocyte response to inflammation, we also performed DEG and pathway analyses for the different time points and treatments. At day1, LPS/nig treatment upregulated pathways, such as NF-kappa B and Th17 signalling, and downregulated pathways related to metabolism and chemokine signalling (Figure 4g; Supplementary Data S2). Notable upregulated genes included *IL6, IL1B,* and *NFKB1,* while *CCL2* was downregulated (Figure 4h; Figure S4g). By contrast, the day 6 cells displayed an upregulation of pathways related to ribosome, Rap1 signalling, and cardiomyopathy in response to LPS/nig treatment, which included genes such as *VEGFA* (Figure 4g,h). Consistent with the cluster analysis, dermal macrophage markers, such as *LYZ* and *TYROBP*, were similarly upregulated in both day 6 conditions (Figure 4h; Figure S4g). These patterns of gene expression further support a model of acute inflammation and monocyte recruitment, followed by resolution and differentiation into tissue resident myeloid cells.

Finally, to explore how monocytes potentially crosstalk with the resident skin cells, we analysed putative receptor-ligand interactions using CellPhoneDB. At day 1, monocytes in the LPS/nig treated models displayed the most predicted interactions with ECs and keratinocytes, while at day 6, most interactions occurred with other monocytes and fibroblasts (Figure 4i). This shift in interactions from keratinocytes to fibroblasts was consistent with early migration of monocytes into the epidermal layer, followed by later migration into the dermis (Figure 2h,i). Further analysis of the significant monocyte-keratinocyte interactions at day 1, revealed potential signalling from the keratinocytes to the monocytes via the prostaglandin E2 and Jagged1-Notch2 pathways and signalling from the monocytes to the keratinocytes via IL-1α and IL-1β pathways (Figure 4j). At day 6, significant signalling interactions from the fibroblasts to the monocytes included ECM-integrin, IL-6, and CSF3 pathways, while monocyte to fibroblast signalling included the VEGF and retinoic acid pathways (Figure 4j). The differentiation into dermal macrophages and cross-talk with fibroblasts at day 6 was further supported by immunofluorescence detection of CD163 positive cells throughout the dermal compartment and close interaction with vimentin positive fibroblasts at this time point (Figure 4k). This analysis therefore demonstrated the ability to uncover putative intercellular regulatory networks within microfluidic HSE and provided new insight how these signalling interactions dynamically change following inflammatory activation.

### Mimicking inflammaging and impaired monocyte responses

Together, the cell tracking and transcriptomic analyses suggested that activation of the microfluidic HSE with LPS/nig promoted an acute inflammatory response, characterised by rapid recruitment of monocytes and upregulation inflammatory cytokines, but after 6 days, the inflammatory response had largely resolved and the remaining monocyte-derived cells differentiated into mature myeloid cells, mostly dermal macrophages. To extend the utility of the model, we next explored whether this response could be perturbed and the microfluidic HSE could be used to mimic a chronic or dysfunctional immune response. Recent studies in human volunteers have shown that there is increased inflammation and monocyte recruitment in the skin of older adults after injection site injuries^21^. Moreover, it has been proposed that this response is mediated by secretion for pro-inflammatory molecules from senescent fibroblasts. We therefore sought to mimic age-associated immune dysfunction with the introduction of senescent fibroblasts in the microfluidic HSEs.

Human dermal fibroblasts were treated with hydrogen peroxide to initiate a stress-induced senescence response, and microfluidic HSEs were then constructed with either untreated proliferative fibroblasts or senescent fibroblasts mimicking young and aged skin, respectively (Figure 5a). Fibroblast senescence was confirmed by increased cell size, upregulation of p21, and loss of Ki67 expression (Figure 5b,c; Figure S5). Proliferative and senescent models were activated with LPS/nig, and monocytes were injected into the vascular channel as before. After 24 hours, immunofluorescence staining for the pan-leukocyte marker CD45 revealed a significant increase in the number of monocytes migrating into the dermal and epidermal compartments of the senescent HSEs stimulated with LPS/nig compared to proliferative HSEs treated with LPS/nig (Figure 5d,e). These findings therefore demonstrated that the microfluidic HSE can also be used to model impaired immune responses associated with ageing and that system accurately replicates in vivo human responses.

**Figure 5:**
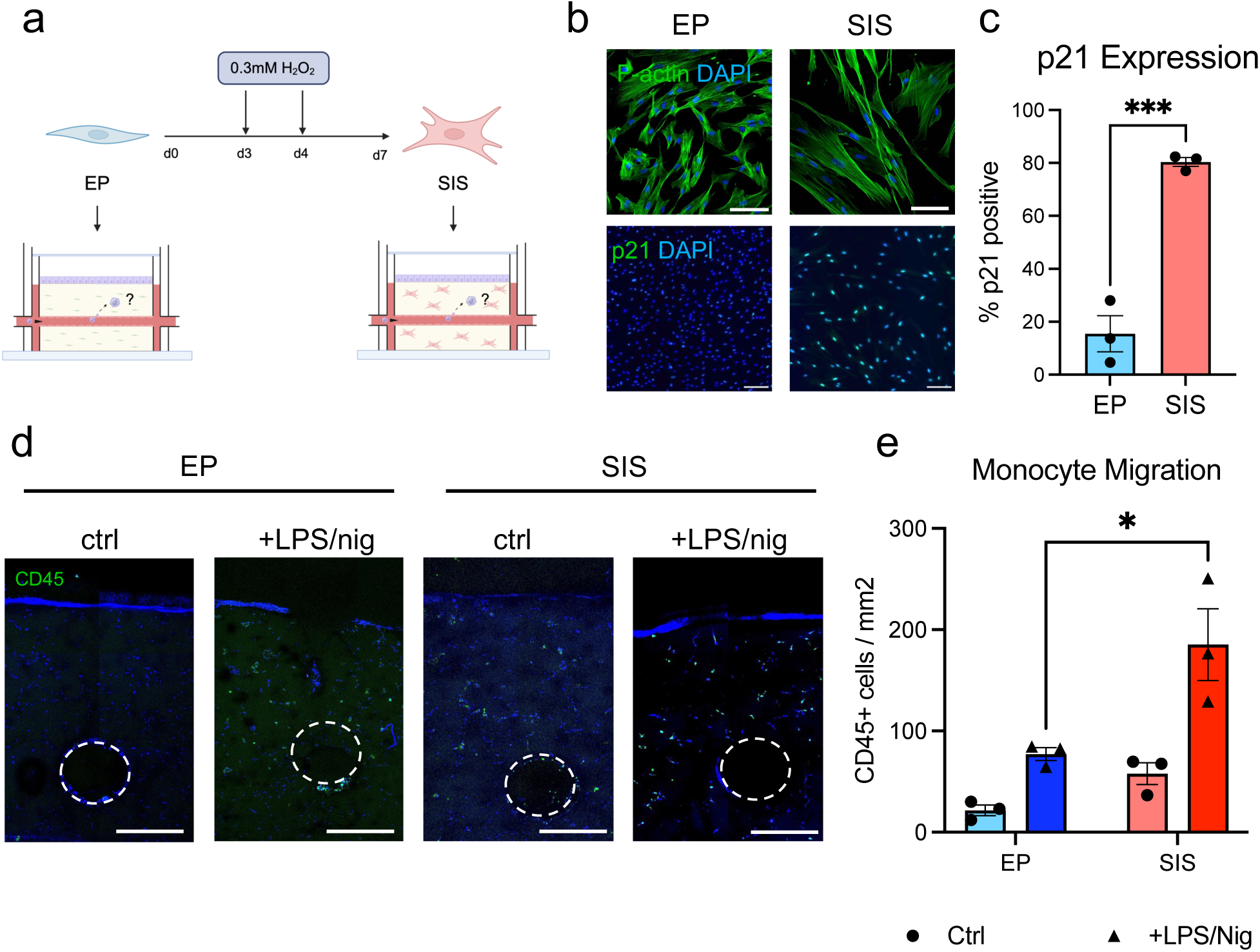
Replicating age-related immune dysfunction with senescent fibroblasts. (a) Schematic of stress-induced senescence protocol and construction of HSEs. Early passage proliferative (EP) human dermal fibroblasts were treated with two doses of 0.3 mM H_2_O_2_ to induce senescence, and HSEs were seeded with EP or SIS fibroblasts in the dermis. HSEs were treated with LPS/nig to initiate inflammation, and monocytes were injected into the microchannel as before. (b) Immunofluorescence images of F-actin and p21 in EP and SIS fibroblasts cultured on glass coverslips for 48 h after completion of the H_2_O_2_ treatment protocol. Scale bar (upper) = 100 μm; scale bar (lower) = 200 μm. (c) Quantification of the percentage of p21 positive cells in EP and SIS fibroblasts on coverslips. Data represent the mean ± SEM of N=3 experiments, ***p<0.001, t-test. (d) Immunofluorescence images of CD45 positive cells (monocytes) in HSEs containing EP or SIS fibroblasts at 24 h after treatment with LPS/nig. Scale bar = 500 μm. (e) Analysis of monocyte migration by quantification of the density of CD45 positive cells outside the microchannel in HSEs containing EP or SIS fibroblasts at 24 h after treatment with LPS/nig. Data represent the mean ± SEM of N=3 experiments, *p<0.05, ANOVA.

## Discussion

In this study we developed and characterised a novel microfluidic HSE model that replicates dynamic immune responses within the skin. Notable features of this system include the ability to deliver circulating immune cells through a vascular microchannel within the dermis, inducible activation of inflammation and monocyte recruitment, and compatibility with live-cell tracking and single-cell transcriptomics. Molecular and cellular characterisation of the microfluidic HSE at single-cell resolution demonstrated that the model accurately replicated key aspects of acute immune responses, such as dynamic up and down regulation of inflammatory mediators in all cell types within the system and differentiation of recruited monocytes into skin resident macrophages. We therefore propose that this platform is well suited for modelling complex and dynamic immune responses in human skin.

A unique advantage of tuneable in vitro models, such as the microfluidic HSE, is the ability to directly manipulate and interrogate the role of specific cell types. By selectively adding or removing cell types, we show here that the keratinocytes are the primary drivers of the inflammatory response to LPS/nig and mediate monocyte recruitment. Secretion of IL-1β and IL-18 by the keratinocytes is consistent with activation of the NLRP3 inflammasome in response to infection or other stressors, as well as its role in stimulating innate immune responses^26–28^. However, we also observed early down regulation of genes associated with inflammation and infection in the keratinocytes, alongside upregulation of protein processing and metabolic pathways. This finding suggests that negative feedback mechanisms in the keratinocytes could help initiate resolution of inflammation, and activation of genes related to unfolded protein response (*DDIT3*) and fatty acid synthesis (*FASN*) could be key players^29,30^. By day 6, expression of inflammatory markers, such as *IL6*, across all cell types was largely resolved, suggesting a return close to baseline, but some notable differences in metabolic genes still persisted. In future studies, this model will be a useful tool for exploring if and over what time scale the system can fully return to baseline, or whether there are persistent effects of inflammatory stress^31,32^.

In addition to acute inflammatory responses, we also demonstrated how the microfluidic HSE can be used to model age-related immune dysfunction. Generation of HSEs with senescent fibroblasts resulted in increased monocyte recruitment into the tissue, consistent with human in vivo studies and the known effects of the senescence-associated secretory phenotype on cytokine production^21^. These data demonstrate that senescent fibroblasts alone are sufficient for over activation of monocyte recruitment, and reports of a similar migratory potential of monocytes from young and older donors^33^, suggest that fibroblasts may be the key drivers of age-related immune dysfunction in the skin. However, further studies are needed to assess the role of distinct monocyte subpopulations^34,35^.

Monocytes within the microfluidic HSE differentiated primarily along a dermal macrophage lineage, and by day 6, most cells expressed markers of tissue resident macrophages, such as *CD163, TYROBP,* and *LYZ*. Although the day 6 monocyte-derived cells also displayed some similarities to DCs and LCs, many of these genes overlap with macrophages, and there was no detectable expression of more definitive DC and LC markers, such as *CD1A* or *CD207*^25^. As monocytes can differentiate into multiple myeloid lineages, it will be interesting to investigate how a wider range of environmental cues and time scales influence differentiation along DC and LC lineages, particularly given the presence of monocyte-derived cells in the epidermis. Moreover, the use of spatial transcriptomics could provide addition insights into the spatial distribution and role of local cues in monocyte differentiation.

While the microfluidic system presented here can replicate several complex features of immune responses within the skin, there are still some limitations. In these studies, monocytes were injected into the microchannel for ease of use rather than delivered by continuous perfusion. As fluid flow is an important regulator of endothelial function^36^, the lack of perfusion could potentially influence monocyte adhesion and trafficking across the endothelium^37^, and a perfusion system could be important to build into next generation platforms. The studies presented here only focused on monocytes as prototypic circulating immune cells, but in the future it will be interesting to investigate a wider range of cell types (e.g. T cells and neutrophils) and to incorporate resident immune cells into the system. However, the addition of adaptive immune cells may require the development of a fully autologous HSE, which would be a significant challenge.

In summary, the microfluidic HSE and associated methodologies represent important advances in our ability to recreate complex immune responses in vitro. The system captures key aspects of immune cell trafficking from the vasculature, inflammatory activation and resolution, and monocyte differentiation into dermal macrophages, and it will be a valuable tool for future investigation into healthy and dysfunctional immune responses within the skin, as well as safety and efficacy testing of therapeutics within a human-based platform.

## Materials and Methods

### 2D Cell culture

N/TERT keratinocytes^38^ were provided by Prof. Michale Philpott’s laboratory and were cultured in complete FAD medium, which comprised: 1 part F12 medium containing sodium pyruvate and GlutaMAX™, 3 parts Dulbecco’s Modified Eagle medium (DMEM) containing sodium pyruvate and GlutaMAX™ (Gibco, ThermoFisher), 10 % fetal bovine serum (FBS; Biosera), 1 % penicillin/streptomycin (PS), 1.8 x 10^-4^ M adenine (Sigma Aldrich), 5 μg/ml insulin (Sigma Aldrich), 0.5 μg/ml hydrocortisone (ThermoFisher), 10 ng/ml mouse epidermal growth factor (Peprotech Inc) and 1 x 10^-10^ M cholera toxin (Sigma Aldrich). N/TERT keratinocytes were passaged at 60-70 % confluency and re-seeded at a minimum density of 0.25 x 10^5^ cells/ml and a maximum density of 1 x 10^5^ cells/ml.

Primary human dermal fibroblasts were obtained from residual tissue of elective abdominoplasty surgeries involving the removal of skin, with written patient consent under the East London and City Health Authority Research Ethics Committee approval (09/HO704/69) and were provided by Prof. Michael Philpott. Fibroblasts were cultured in DMEM containing sodium pyruvate and GlutaMA (Gibco, ThermoFisher) and supplemented with 10 % FBS and 1 % PS. Fibroblasts were passaged once they reached 70-80 % confluency and were re-seeded at a minimum density of 2.5 x 10^4^ cells/ml, up to passage 10. To induce senescence, fibroblasts were treated with 0.3 mM H_2_O_2_ in normal growth medium on two sequential days and then cultured for three additional days before seeding into microfluidic HSEs.

ECs (C2519A, LONZA) were cultured in Endothelial Growth Medium 2 (EGM2) containing SupplementMix (PromoCell) and 1 % PS. The final concentration of the components within SupplementMix was: 2 % FBS, 5 ng/ml recombinant human epidermal growth factor, 10 ng/ml recombinant human basic fibroblast growth factor, 20 ng /ml recombinant human insulin-like growth factor I (IGF-I), 0.5 ng/ml recombinant human vascular endothelial growth factor (VEGF), 1 µg/ml ascorbic acid, 22.5 μg/ml heparin, and 0.2 µg/ml hydrocortisone. Cells were passaged at 90 % confluency and were cultured to a maximum passage number of 5.

THP-1 cells (TIB-20, ATCC) were cultured in suspension in RPMI-1640 culture medium supplemented with L-glutamine (ThermoFisher), 10 % FBS and 1 % PS. THP1 cells were cultured at a minimum density of 2 × 10^5^ cells/ml and were passaged when reaching 8 × 10^5^ cells/ml. All cells were cultured at 37 °C, 5 % CO_2_.

### Monocyte isolation

Ethical approval was provided by the Queen Mary Ethics of Research Committee, reference number QMERC23.059. Individuals 18 years or older were recruited into the study, and full written informed consent was obtained from each donor. Up to 80 ml of peripheral blood was taken from each donor and processed as described below. Individuals were excluded from the study if they had a recent history (≤5 years) of neoplasia, immunosuppressive disorders which required immunosuppressive medication, were anaemic, or had a current pregnancy. Peripheral blood mononuclear cells were isolated by density centrifugation using Ficoll-Paque (Amersham Biosciences). Monocytes were isolated by positive selection using the CD14+ positive isolation kit according to the manufacturer’s instructions (Miltenyi Biotec).

### 3D printing of microfluidic skin equivalents

Microfluidic HSEs were constructed using the 3D Discovery Bioprinter equipped with a class II biosafety cabinet (RegenHU) and an extrusion-based print head. The BioCAD software (RegenHU) was used to create the layer-by-layer designs of the custom frame and microchannel template.

Silicone Elastomer 1700 (SE1700, DOWSIL, Corning) was used to print custom frames for the microfluidic HSEs. The SE1700 was prepared by mixing the base with the catalyst at a 10:1 ratio and was printed at a 5 mm/s feed rate, using a 27 G needle (Metcal, OK International). The outer frame measured 24 mm by 24 mm and was printed to have a build height of 9.6 mm. The compartment for the media measured 20 mm by 20 mm and was printed with a build height of 8 mm. The internal compartment, which contained the 3D microfluidic model, was 15 mm by 10 mm and was built to a height of 2 mm. The frames had an inlet and outlet for access to the microchannel, within the HSEs, for the injection of cells. The inlet and outlets were 0.8 mm in height and 2 mm wide. After printing, the frames were cured at 70 °C for 4 h and autoclaved before sterile use.

For the generation of the dermal compartment, 25 μl of a 1 x 10^6^ cell/ml cell suspension of fibroblasts (in complete DMEM) was combined with 75 μl of sterile 10 U/ml thrombin (Sigma Aldrich) in Dulbecco’s phosphate buffered saline (DPBS) (Sigma Aldrich) containing 0.1 % bovine serum albumin (BSA) (ThermoFisher) and 100 μl of sterile 40 mg/ml fibrinogen from bovine plasma (Sigma Aldrich) in complete EGM2. The solution was immediately pipetted into the inner compartment of the custom frame, to create a fibroblast-embedded fibrin base layer.

7 % (w/v) (gelatin type A from porcine skin) in DPBS supplemented with 1 % PS was printed directly onto the fibrin base layer, to create a sacrificial microchannel template. The gelatin ink was printed at 26 °C, using a 32 G needle (Metcal, OK International) and a 1 mm/s feed rate. The constructs were then incubated at 4 °C in the fridge for 30 min to solidify the gelatin. The microchannel template was then embedded in another layer of fibroblast-embedded fibrin.

The microchannel template was selectively removed by incubating the construct at 37 °C for 1 h, to melt the gelatin. DPBS, warmed to 37 °C, was then flushed through the microchannel 3 times, using the inlet and outlet of the frame, to remove the melted gelatin.

A 1 x 10^7^ cells/ml EC cell suspension was prepared in complete EGM2 medium. 1.5 x 10^4^ ECs were injected into the microchannel (approximately 1.5 μl). The dermal compartment was then turned upside down and incubated at 37 °C for 45 min. After 45 min, the dermal compartment was returned to normal orientation and EC attachment to the upper lining of the microchannel was confirmed using brightfield microscopy. 1.5 ml of complete EGM2 medium was then pipetted on top of the dermal compartment and the constructs were cultured for 72 h, to allow the formation of a confluent endothelial lining of the microchannel.

5 x 10^5^ N/TERT keratinocytes suspended in 100 μl of FAD were seeded on top of the construct and the microfluidic HSE was incubated at 37 °C for 1 h to allow keratinocyte attachment. After 1 h, the construct was washed twice with 1.5 ml DPBS to removed unattached cells. 1.5 ml of reduced hydrocortisone (0.1 μg/ml) FAD medium was then pipetted on top of the construct. The microfluidic HSE was then cultured for 24 h.

### Induction of inflammation

Microfluidic HSEs were treated with 500 ng/ml of LPS (Cell Signaling Technology) in 1.5 ml of complete EGM2 medium for 4 h. Microfluidic HSEs were then treated with 10 μM nigericin (Tocris Bioscience) in 1.5 ml of complete EGM2 medium for 1 h. THP-1 or primary CD14+ monocytes were suspended in complete RPMI medium at 7 x 10^6^ /ml and 1.05 x 10^4^ cells were injected into the microchannel of each HSE (approximately 1.5 μl). The microfluidic HSEs were then cultured for 3 h, 24 h, or 6 days. For the day 6 cultures, the media was replaced every 2 days.

### Membrane labelling and immunofluorescence staining

CellBrite cytoplasmic membrane labelling kits (Green, Orange, or Red; Biotium) were used to fluorescently label the cytoplasmic membranes with cells within the microfluidic HSEs. For staining, cells were suspended at a density of 1×10^6^ /ml in normal growth medium. 5 μl of CellBrite dye was added per 1 ml of cell suspension. The cell suspensions were mixed well and incubated for 30 min at 37 °C. After 30 min the cells were centrifuged at 300 G for 5 min. The supernatant was discarded and the cells were washed once, by resuspending the cells in normal growth medium at 3x the staining volume. The cells were centrifuged again at 300 G for 5 min and resuspended in normal cell culture media, at the desired seeding concentration.

Microfluidic HSEs were fixed in 4% paraformaldehyde (PFA) for 15 min and washed 3 times with DPBS. Samples were cut transversely through the microchannel into 4 pieces, and each slice was permeabilised using 0.05 % Triton-X 100 (Sigma Aldrich) in DPBS for 15 min, at room temperature. The samples were then washed with DPBS and blocked for 2 h using 10 % FBS and 0.25 % fish gelatin (Sigma Aldrich) in DPBS. Samples were then incubated at 4 °C with primary antibodies for either keratin 14 (1:50 dilution, mouse, LL002, Ab7800 Abcam), vimentin (1:50 dilution, rabbit, D21H3, 5741T Cell Signaling), CD31 (1:50 dilution, mouse, WM59, MCA1738 BioRad), CD68 (1:50 dilution, mouse, KP1 916104 Biolegend), CD163 (1:25 dilution, directly conjugated, mouse, GHI/61, 333620 Biolegend) or CD45 (mouse, HI30, 304001 Biolegend) in the blocking solution, for 48 h.

Samples were washed 3 times with DPBS before being incubated with secondary antibodies and 4′,6-diamidino-2-phenylindole (DAPI, Invitrogen, 10125092) at 1:500 in the blocking solution, for 24 h, at 4 °C. Secondary antibodies were either goat anti-mouse (Invitrogen, A-11001, Alexa Fluor 488), donkey anti-mouse (Invitrogen, A-11037, Alexa Fluor 568), goat anti-rabbit (Invitrogen, A-21428, Alexa Fluor 555) or goat anti-rabbit (Invitrogen, A-11008, Alexa Fluor 488). Samples were washed 3 times with DPBS and transferred into custom printed SE1700 staining frames on microscope slides (Gerhard Menzel GmbH). 100 μl of DPBS was added on top of the samples and 24 x 24 mm coverslips (Epredia Menzel Microscope Coverslips) were mounted on top using vacuum grease (ThermoFisher). Samples were stored at 4 °C if not immediately imaged.

The LSM 880 Confocal Microscope with Airyscan (Zeiss) was used for either live imaging of fluorescently tagged cells within the microfluidic HSEs or imaging of fixed immunostained samples, and ImageJ was used for image analysis. For quantification of monocyte migration in living cultures or CD68 stained samples, Z-stack images were acquired through the full thickness of the HSE, and sub-stacks for the epidermal layer or only the dermal compartment above the microchannel were used to generate maximum intensity projections. The number of monocytes were then detected by particle analysis in the 2D projected image and reported as cells per unit area. For quantification of CD45+ cell migration, cross-sections of HSE slices were imaged by confocal microscopy, and the total number of cells outside the channel were detected by particle analysis and reported as cells per unit area.

### Cytokine analysis

The Proteome Profiler Human Cytokine Array (R&D Systems) was used to detect 36 human cytokines, chemokines and acute phase proteins simultaneously from the conditioned medium of untreated control keratinocytes or keratinocytes treated with LPS/nig 24 h prior. Membranes were prepared following the manufacturer’s instructions and incubated with 1 ml of Chemi Reagent Mix (R&D Systems) for 1 min, before being visualised and imaged on the ChemiDoc XRS+ (BioRad). The ImageLab software (BioRad) was used to determine the average signal (pixel density) of the pair of duplicate spots representing each cytokine.

### Single-cell RNA sequencing

Three microfluidic HSEs containing primary CD14+ monocytes were generated per condition (Ctrl Day1, LPS/nig Day1, Ctrl Day6, and LPS/nig Day6), using the methods described above. For single-cell RNA sequencing (scRNA-seq), the samples were removed from the frames and digested with 300 μl of 50 FU/ml nattokinase (MedChem Express) dissolved in sterile DPBS for 1 h at 37 °C 5 % CO_2_. Cell suspensions from each microfluidic HSE were transferred into sterile 15 ml tubes and centrifuged at 200 G for 15 min at 4 °C. The supernatant was discarded and cell pellets from each sample were resuspended in 40 μl of FACs buffer and kept on ice.

A CD45 positive selection was performed according to the manufacturer’s instructions (Miltenyi Biotec), to separate monocyte-derived cells from fibroblasts, ECs, and keratinocytes. The CD45 positive and negative samples were resuspended in 500 μl DPBS and filtered through 70 µm pore sterile Flowmi Cell Strainers (Merck, Sigma Aldrich). Acridine orange and propidium iodide (Allied Genetics Inc) were used to assess the viability of single cell suspensions according to the manufacturer’s instructions, and a viability cut-off of > 80 % was set for each sample to proceed. 7,500 live CD45 negative cells were combined with 7,500 CD45 positive live cells, from each condition, and kept on ice.

Single-cell RNA sequencing (scRNA-seq) was performed by the Genome Centre, Queen Mary University of London using the Chromium Next GEM Single Cell 3’ assay (10X Genomics). Libraries (3’ v3.1) were generated according to the manufacturer’s instructions, and library QC was performed using the Qubit 2.0 Fluorometer (ThermoFisher) and the Agilent 4200 Tapestation. Final libraries were sequenced using the Illumina NextSeq2000 system to achieve a minimum depth of 50,000 raw reads per cell. NextSeq2000 QC resulted in a quality score (% bases > Q30) of 94.91 %. 15,000 cells were loaded per lane (4 lanes, 15,000 cells per condition) of the Single Cell chip (10X Genomics). A total of 35,987 cells were sequenced (14,410 cells from the Control Day 1 condition, 7516 cells from the Treated Day 1 condition, 7999 cells from the Control Day 6 condition and 6062 cells from the Treated Day 6 condition). FASTQ files were generated using the 10X Genomics Cell Ranger pipeline and empty droplet removal was performed using the EmptyDrops method incorporated in the Cell Ranger software.

### Transcriptomic data analysis

ScRNA-seq data were analysed using the Seurat toolkit (v 5.0.1) in RStudio. Cell Ranger files (features, barcodes, and matrix) were read and converted into a Seurat object and then filtered for cells with unique feature counts above 200, total counts below 50,000, and mitochondrial counts below 15%. Data sets for the different treatment and time points were integrated using the Seurat v5 integration procedure, and the standard workflow was used to normalise, find variable features, and scale the data. The integrated data set was clustered using the Louvain algorithm with 18 principal components (PCs) and resolution of 1 and visualised by UMAP dimensional reduction. The number of PCs and resolution in the initial analysis were adjusted such that there were a minimum number of clusters while each aligned to one of the four major cell types in the model. Cell type identification for the keratinocytes, fibroblasts, ECs, and monocytes was performed using canonical markers for each cell type (keratinocytes: KRT14, KRT5, DSP, LAMA3; fibroblasts: PDGFRA, COL1A1, COL1A2, COL6A3; monocytes: FCER1G, CCL3, CD14, PTPRC; endothelial cells: PECAM1, ECSCR, CLDN5, HSPG2) and visualised using the DotPlot command. Analysis of differentially expressed genes between treatments and time points for each cell type was performed using the FindAllMarkers command on subsets for each cell type, grouped by condition. Differentially expressed genes were identified based on Log2FC > 1, P_adj_ < 0.05, and percentage of expressing cells >10%. Subsequent analysis of up and down regulated pathways by LPS/nig treatment at days 1 and 6 was performed using Gene Set Enrichment Analysis (WebGestalt; KEGG database). Patterns of differentially expressed genes were plotted using DoHeatMap, FeaturePlot, and VlnPlot in Seurat.

Additional analysis of the monocyte populations was carried out by re-clustering the monocyte subset using 15 principal components and resolution of 0.7 and visualisation by UMAP dimensional reduction. Markers for the eight resulting monocyte clusters were identified using the FindAllMarkers command and then compared to conserved markers for all myeloid populations in human skin^25^. A relative similarity score was calculated by the percentage of the conserved in vivo markers found in the top 100 markers for each monocyte-derived in vitro cluster. Trajectory analysis of the monocytes was performed using the Monocle3 toolkit in RStudio, and calculated pseudotime values were replotted as a feature in the Seurat UMAP plot. Predicted intercellular interactions were analysed using the CellPhoneDB toolkit in Python. The statistical inference method was used to determine significant interactions between the four major cell types for each treatment and time point. Total interactions with monocytes and selected molecular interactions were plotted with ktplots in R using the heatmap and dot plot functions, respectively.

### Statistical analysis

GraphPad Prism was used to perform statistical analysis. Acquired data were analysed using either unpaired t-tests, one-way or two-way analysis of variance (ANOVA) with Tukey multiple comparisons test. When calculating overall means from independent experimental repeats, the results were reported as mean ± standard error (SEM). A p-value < 0.05 was considered significant.

## Acknowledgements

This study was funded by a PhD studentship from the Medical Research Council and a project grant from the Biotechnology and Biological Sciences Research Council (BB/X006972/1). Dr Chambers was funded by Barts Charity (MGU0459). We thank Prof. Clare Bennett for insight and guidance on the analysis of monocyte fate and Prof. Cleo Bishop for assistance with the induction and analysis of fibroblast senescence.

## Supplementary Figures

**Figure S1:**
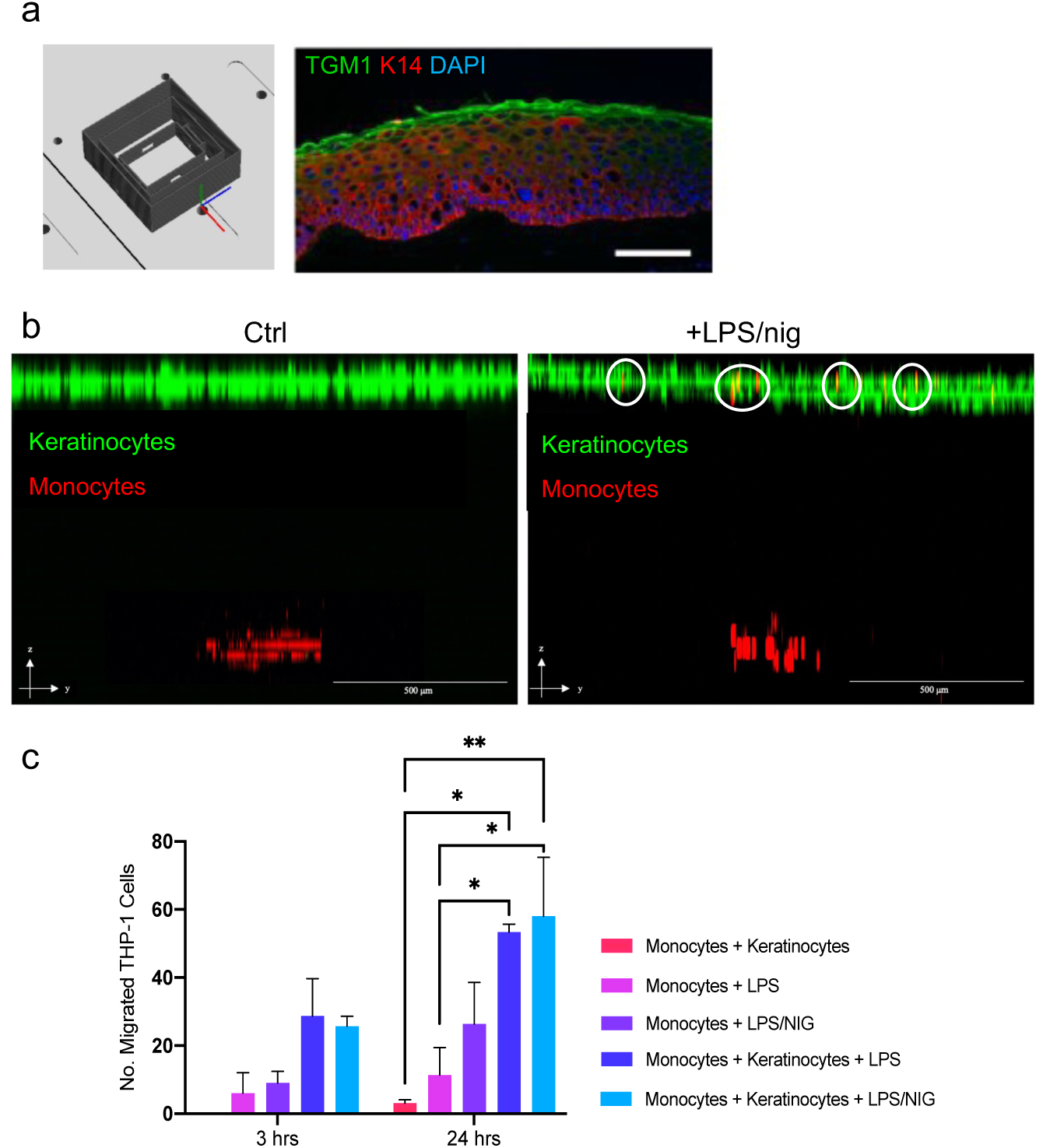
Development of microfluidic HSE and analysis of monocyte migration. (a) 3D CAD design of chamber for air-liquid interface cultures and immunofluorescence image of K14 and tranglutaminase-1 in stratified epidermal layers of HSEs at day 14. (b) Cross-sectional views of confocal Z-stacks for HSEs containing fluorescently labelled keratinocytes and THP-1 monocytes, 24 h after control or LPS/nig treatment. (c) Quantification of monocytes per field of view in the epidermis in HSEs with monocytes alone or monocytes and keratinocytes at 3h and 24 h after control, LPS, or LPS/nig treatment. Data represent mean ± SEM of N=3 experiments, *p<0.05, **p<0.1, ANOVA.

**Figure S2:**
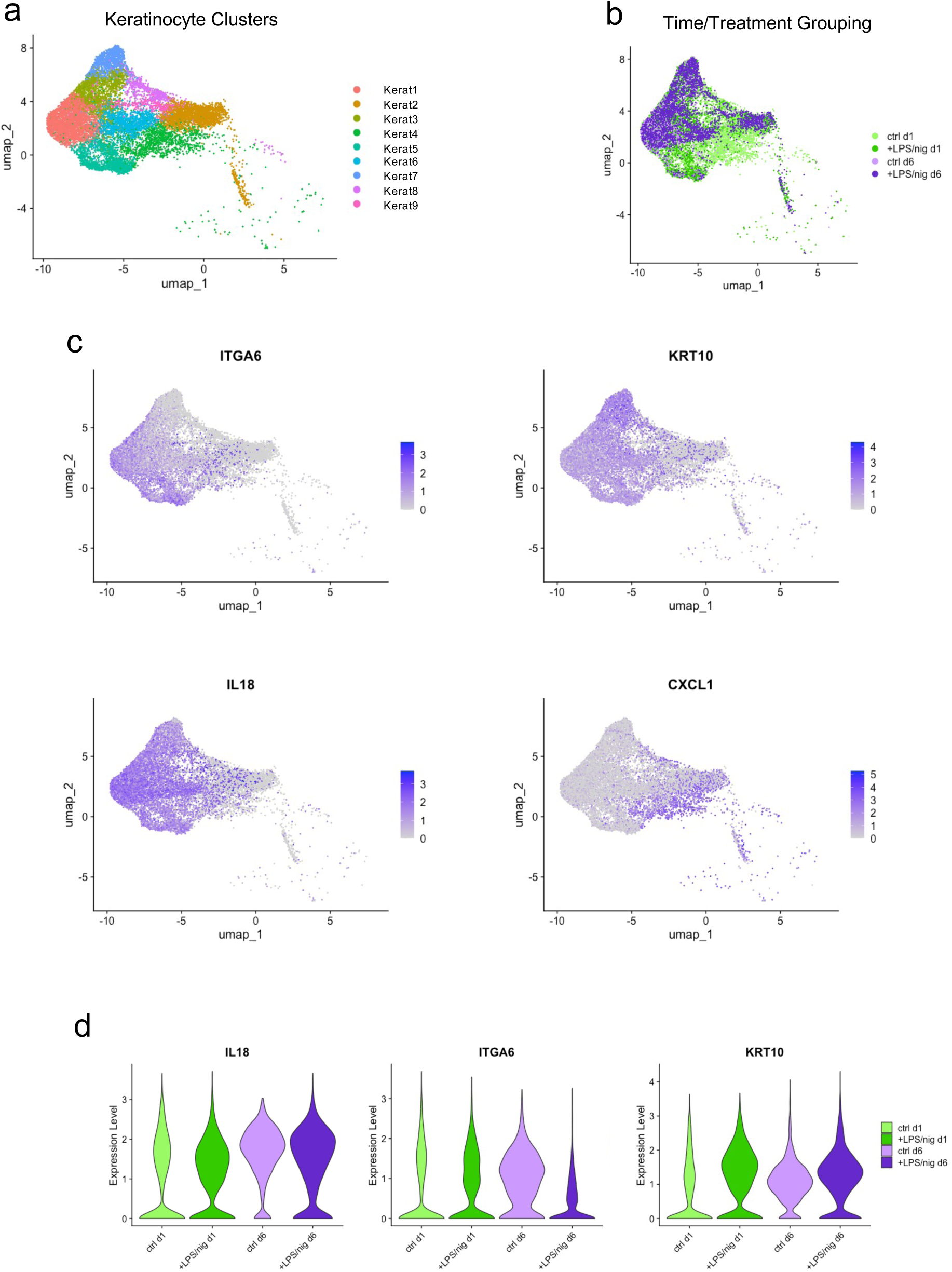
Analysis of keratinocyte clusters. (a) UMAP visualisation of all keratinocyte clusters in the single-cell RNA-seq data set. (b) Feature plot of time point and treatment conditions in keratinocyte clusters. (c) Feature plots for key discriminatory markers for keratinocytes clusters, including differentiation (*ITGA6, KRT10*) and inflammatory genes (*IL18, CXCL1*). (d) Violin plots of *IL18, ITGA6,* and *KRT10* across the experimental conditions.

**Figure S3:**
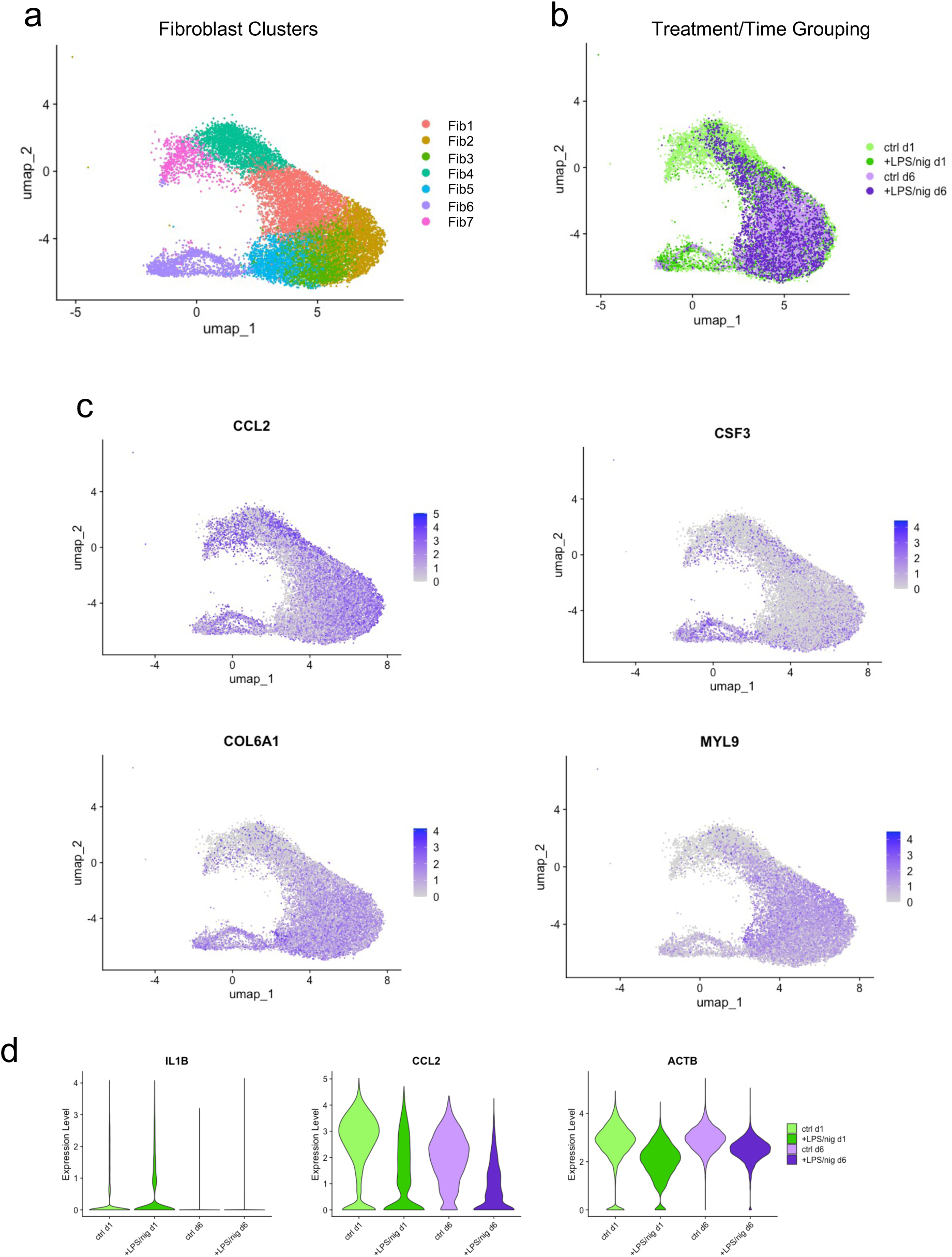
Analysis of fibroblast clusters. (a) UMAP visualisation of all fibroblast clusters in the single-cell RNA-seq data set. (b) Feature plot of time point and treatment conditions in fibroblast clusters. (c) Feature plots for key discriminatory markers for fibroblast clusters, including *CCL2, CSF3, COL6A1, MYL9*. (d) Violin plots of *IL1B, CCL2,* and *ACTB* across the experimental conditions.

**Figure S4:**
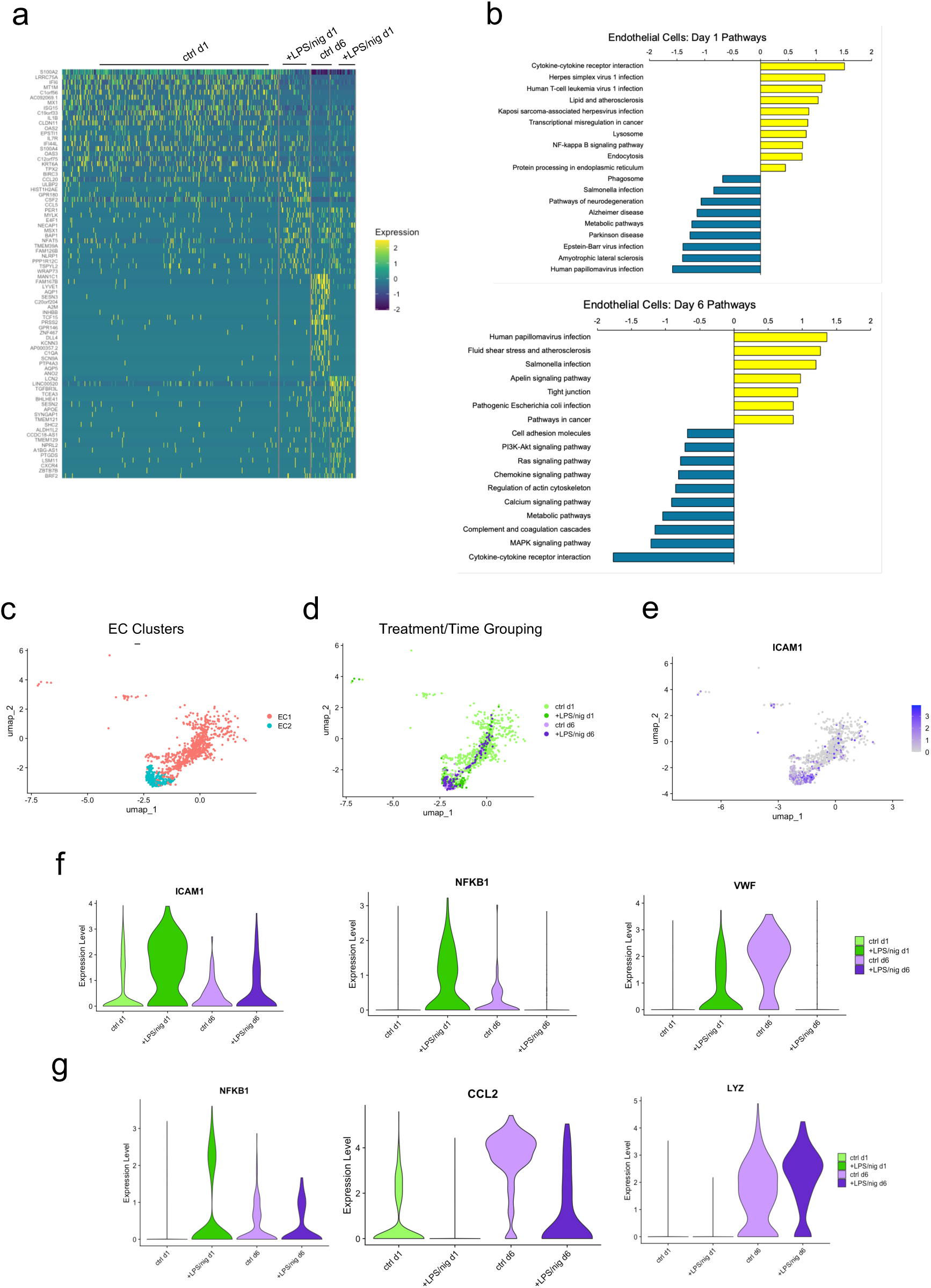
Analysis of EC clusters and pathways and additional monocyte genes. (a) Heatmap of the top 20 differentially expressed genes (DEGs; Log2FC > 1; Padj < 0.05) between experimental conditions for all EC clusters. (b) Gene set enrichment analysis (GSEA; Webgestalt) of EC pathways up and down regulated by LPS/nig at days 1 and 6. Data represent pathways with the top 10 highest and lowest relative enrichment scores. (c) UMAP visualisation of the two EC clusters in the single-cell RNA-seq data set. (d) Feature plot of time point and treatment conditions in EC clusters. (e) Feature plots for *ICAM1*. (f) Violin plots of *ICAM1, NFKB1, and VWF* across the experimental conditions. (g) Violin plots of *NFKB1*, *CCL2*, and *LYZ* in monocytes.

**Figure S5:**
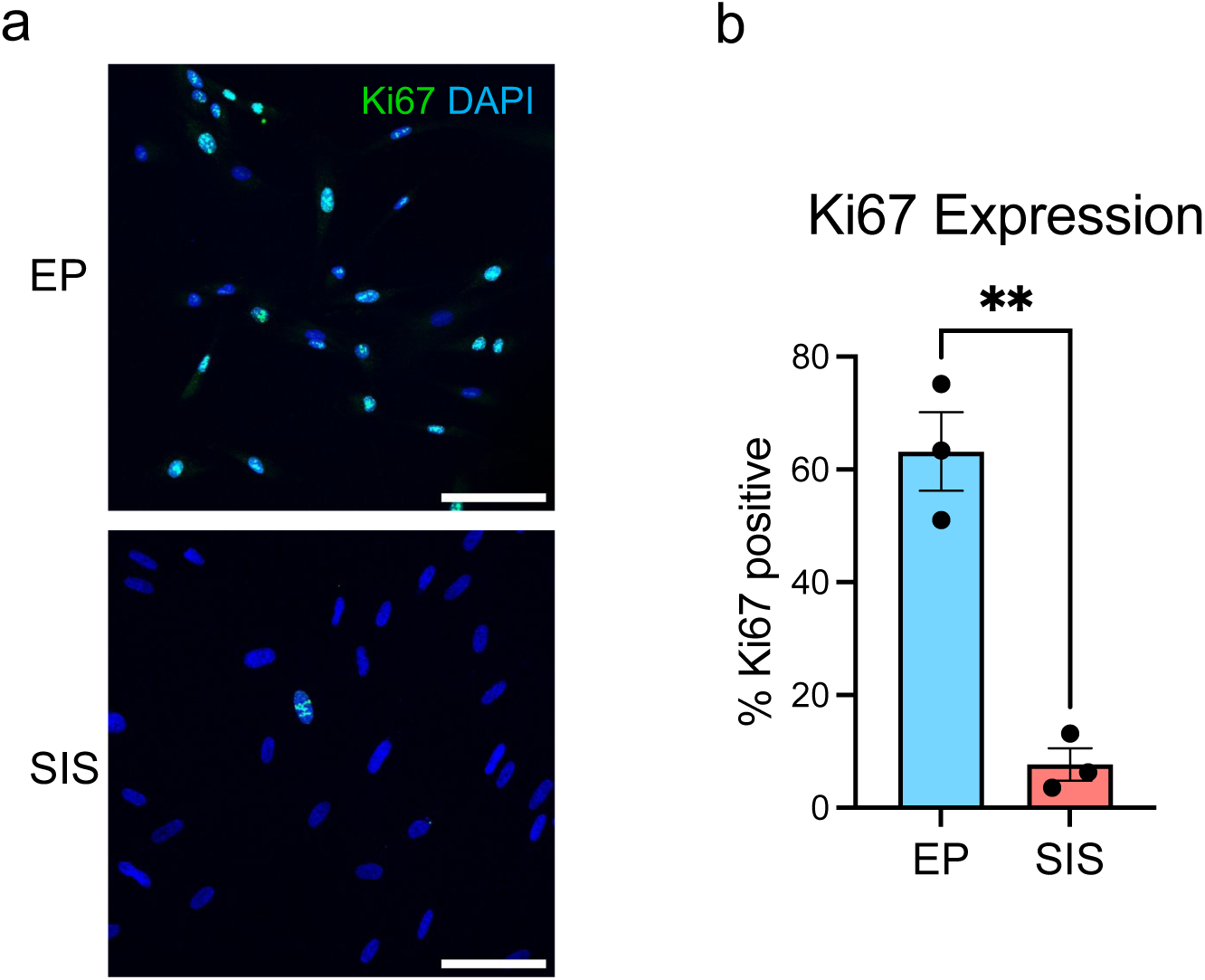
Validation of cell cycle arrest in senescent fibroblasts. (a) Immunofluorescence images of Ki67 in EP and SIS fibroblasts cultured on glass coverslips for 48 h after completion of the H_2_O_2_ treatment protocol. Scale bar = 100 μm. (c) Quantification of the percentage of Ki67 positive cells in EP and SIS fibroblasts on coverslips. Data represent the mean ± SEM of N=3 experiments, **p<0.005, t-test.

